# Precision-dependent modulation of social attention

**DOI:** 10.1101/2024.09.17.612568

**Authors:** Wenhui Gao, Changbo Zhu, Bailu Si, Liqin Zhou, Ke Zhou

## Abstract

Social attention, guided by cues like gaze direction, is crucial for effective social interactions. However, how dynamic environmental context modulates this process remains unclear. Integrating a hierarchical Bayesian model with fMRI, this study investigated how individuals adjusted attention based on the predictions about cue validity (CV). Thirty-three participants performed a modified Posner location-cueing task with varying CV. Behaviorally, individuals’ allocation of social attention was finely tuned to the precision (inverse variance) of CV predictions, with the predictions being updated by precision-weighted prediction errors (PEs) about the occurrence of target locations. Neuroimaging results revealed that the interaction between allocation of social attention and CV influenced activity in regions involved in spatial attention and/or social perception, such as the temporoparietal junction (TPJ), frontal eye field (FEF), superior temporal sulcus (STS), and inferior parietal sulcus (IPS). Precision-weighted PEs about target locations specifically modulated activity in the TPJ, STS, and primary visual cortex (V1), underscoring their roles in refining attentional predictions. Dynamic causal modeling (DCM) further demonstrated that enhanced absolute precision-weighted PEs about target locations strengthened the effective connectivity from V1 and STS to TPJ, emphasizing their roles in conveying residual error signals from low-level sensory areas to high-level critical attention areas. These findings elucidated how the precision of contextual predictions dynamically modulated social attention, offering insights into the computational and neurocognitive mechanisms of context-dependent social attention.

## Introduction

In everyday social interactions, our gaze naturally follows others’ eye movements or pointing gestures, a process known as social attention. Social attention, triggered by cues, such as gaze direction, head orientation, and body movements (Kleinke, 1986; Lawson & Calder, 2016), is essential for understanding others’ intentions and facilitating effective communication (Behrens et al., 2009). This evolutionarily adaptive mechanism enhances our ability to navigate social environments. To study social attention, researchers often use paradigms like Posner’s location-cueing task (Posner, 1980; Posner et al., 1980), which demonstrates that responses are faster when targets appear at cued locations compared to uncued ones, reflecting an orientating of covert attention. However, the neurocognitive mechanisms underlying this process remained complex and not fully understood, involving ongoing debates about whether it is purely reflexive (i.e., exogenous), voluntary (i.e., endogenous), a combination of both, or a distinct type of attention.

Previous research has shown that participants respond faster to targets appearing at gaze-cued locations than at uncued locations, regardless of whether gaze reliably predicts target location or even counter-predicts target location (Driver et al., 1999; Friesen et al., 2004; Friesen & Kingstone, 1998), implying a reflexive component in social attention. However, the magnitude of the gaze-cueing effect increased with cue validity (CV, the percentage of cue being valid to target) (Hill et al., 2010) and correlated with self-reported voluntary control (Tipples, 2008), indicating a voluntary component in social attention. While both social attention and endogenous attention exhibit slow attentional facilitation, only social attention is followed by an inhibition of return (IOR) effect (Müller & Findlay, 1988; Frischen & Tipper, 2004), typically found in exogenous attention triggered by peripheral cues. However, peripheral cues usually induce IOR within 100 ms of cue onset (Posner et al., 1985), whereas gaze cues induce IOR with a delayed onset compared to peripheral cues (Frischen & Tipper, 2004). Together, these findings suggest that social attention may involve distinct mechanisms from those of purely exogenous and endogenous attention.

Previous neurophysiological studies have continued to fuel the ongoing debate regarding the relationship between gaze-cued social attention and arrow-cued non-social attention, extending the discussion from behavioral evidence to neural correlates. Some research has identified overlapping neural substrates for both attention types, with shared activations in the superior temporal sulcus (STS), inferior parietal lobule (IPL), inferior frontal gyrus (IFG), and the occipital areas (Sato et al., 2009). Common activations in the STS and IFG have been observed within 200-400 ms after cue onset (Uono et al., 2014). However, other studies suggested distinct neural correlates for each attention type. For instance, Hietanen et al. (2006) identified regions where the activation strength was greater for arrow cues than for gaze cues, including the inferior occipital gyrus (IOG), medial temporal gyrus (MTG), precuneus, frontal eye field (FEF), and supplementary eye field (SEF). In contrast, studies using ambiguous cues have reported a greater blood oxygenation level-dependent (BOLD) response in regions like the middle occipital gyrus (MOG), middle frontal gyrus (MFG), precentral gyrus (PreCG), and right STS (rSTS) when the cue was perceived as eye gaze than as arrow (Kingstone et al., 2004; Tipper et al., 2008). Additionally, Lockhofen et al. (2014) demonstrated gaze > arrow activation in the superior occipital gyrus (SOG) and fusiform gyrus (FFG), and arrow > gaze activation in the MOG and right superior parietal gyrus (rSPG). Moreover, studies comparing invalid to valid cues indicated that invalid gaze cues enhanced activation in the temporo-parietal junction (TPJ) and IPL, regions associated with the ventral attentional network (VAN), as well as the STS and FFG, regions involved in face processing; conversely, arrow cues uniquely engaged regions in the dorsal attentional network (DAN) (Engell et al., 2010; Joseph et al., 2015). Additionally, Tipper et al., (2008) reported higher P1 amplitudes for cued versus uncued targets in gaze-induced attention, but not in arrow-induced attention. These findings collectively suggested that social attention uniquely engages face processing regions and enhances bottom-up sensory gain more than top-down control, compared to endogenous attention.

The neural mechanisms of social attention also differ from exogenous attention. Electroencephalography (EEG) studies showed that only valid peripheral cues elicited larger early-stage occipital P1/N1 components than invalid ones, while only invalid gaze cues produced a larger P3 component compared to valid ones, reflecting greater stimulus enhancement in exogenous attention and an influence on later decision-making processes in social attention (Chanon & Hopfinger, 2011). Moreover, patients with frontal-lobe lesions exhibited impaired social attention but retained intact exogenous attention (Vecera & Rizzo, 2006). The findings suggested that social attention enhances top-down control more than bottom-up sensory gain compared to exogenous attention.

Traditional neurophysiological studies have identified brain regions involved in social attention, but have yet to clarify the mechanisms driving their involvement. Prior research, largely based on qualitative contrasts, has provided fragmented insights. Computational modeling provides a quantitative framework to infer cognitive mechanisms underlying social attention, addressing limitations of qualitative methods. This approach can further integrate behavioral data with neurophysiological evidence, advancing our understanding of the social attention process.

The current study aimed to explore the neural mechanisms underlying social attention by applying computational modeling techniques within a hierarchical Bayesian framework. This framework posited that individuals continuously generate and refine hierarchical predictions based on dynamic environmental statistics (Dayan et al., 1995; Friston, 2009; Mathys et al., 2011, 2014). Previous research has successfully applied this approach to examine voluntary spatial attention triggered by arrow cues in the Posner location-cueing paradigm, revealing that attention orientation was modulated by the precision (inverse variance) of CV predictions over time (Vossel et al., 2014a, 2014b). Corresponding neural activities in the TPJ, FEF, and putamen aligned with this precision-dependent attention (Vossel et al., 2015; Kuhns et al., 2017). Additionally, within the hierarchical Bayesian framework, CV predictions were updated by precision-weighted prediction errors (PEs) about target locations (Mathys et al., 2011; Vossel et al., 2014a). Subsequent research has demonstrated that these precision-weighted PEs modulate activities across the frontal gyri, anterior cingulate cortex (ACC), inferior parietal sulcus (IPS), anterior insula, and dopaminergic midbrain areas (Diaconescu et al., 2017; Iglesias et al., 2013, 2021).

This study extended the hierarchical Bayesian model to examine the dynamics of social attention, specifically through eye gaze cues. By combining fMRI data with Bayesian modeling of behavioral data, we aimed to elucidate the computational neural mechanisms underlying social attentional allocation and assess the precision-dependent updates in attention shifts. Additionally, we explored the neural correlates of predictions and PEs. Finally, dynamical causal modeling (DCM) was utilized to analyze how these predictions and PEs modulate information transfer across the involved neural networks.

## Materials and methods

### Participants

To determine the required sample size for the main study, a preliminary pilot study was conducted with 7 participants (4 males, 3 females; median age 22, range 19–25). Sample size estimation was performed using G*Power (version 3.1.9.7; Faul et al., 2009), based on the effect size 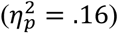 and the average correlation among repeated measures (*r* = .70) derived from the pilot study. The analysis indicated that, to achieve a statistical power of .95 with an αlevel of .05, a minimum sample size of 7 participants was needed. To ensure the robustness of the findings, a sample size larger than this minimum requirement was utilized in the main study.

Forty-six healthy volunteers, who had signed informed consent in advance, participated in the present experiment. To ensure optimal task engagement, stringent inclusion criteria were applied. Participants with behavioral accuracy lower than 97% (N = 9 participants), head movement in fMRI exceeding 3 mm (N = 3), or who reported physical discomfort during MR scanning (N = 1) were excluded from further analysis. This resulted in a final sample of 33 participants (9 males, 24 females; median age 21, range 19–24), all of whom were right-handed, had normal or corrected-to-normal vision, and had no history of neurological or psychiatric conditions. The current study was approved by the Institutional Review Board of Beijing Normal University.

### Experimental paradigm

All stimuli were generated and presented using MATLAB (2018b, MathWorks) with the Psychophysics Toolbox Version 3 extension (PTB-3; Kleiner et al., 2007) on a 24-inch monitor (spatial resolution 1920 ×1080 pixels, 60-Hz refresh rate) at the back of the magnet bore. Participants viewed the stimuli through a mirror attached to the head coil, with a viewing distance of 83 cm.

A modified version of Posner’s location-cueing paradigm (Posner, 1980; Sevgi et al., 2020) was applied, as depicted in Fig. 1A. Each trial started with a central green fixation cross (0.5° wide of the visual angle) on a gray background. After a fixation period of 2000 ms (±500 ms jitter), a face with eyes looking straight ahead (5.4° ×6.1°) appeared at the screen center for 100 ms, followed by a rapid gaze shift to the left or right, serving as the social cue for 400 ms. After a blank of 100 ms, a peripheral circular grating (1.5° radius) appeared at 7° eccentricity to the left or right of the fixation cross for 100 ms. In catch trials, the grating did not appear. Trials where the grating appeared in the same direction as the social cue pointed were valid trials, while those where it appeared in the opposite direction were called invalid trials. The gratings were randomly oriented either vertically or horizontally to minimize stimulus adaptation effects. Participants were instructed to maintain their gaze on the central fixation cross throughout the trial and to respond as quickly and accurately as possible by pressing a button if they detected the appearance of a grating within a 1500 ms response window. Each participant completed two sessions, each lasting approximately 20 minutes.

**Fig. 1.**
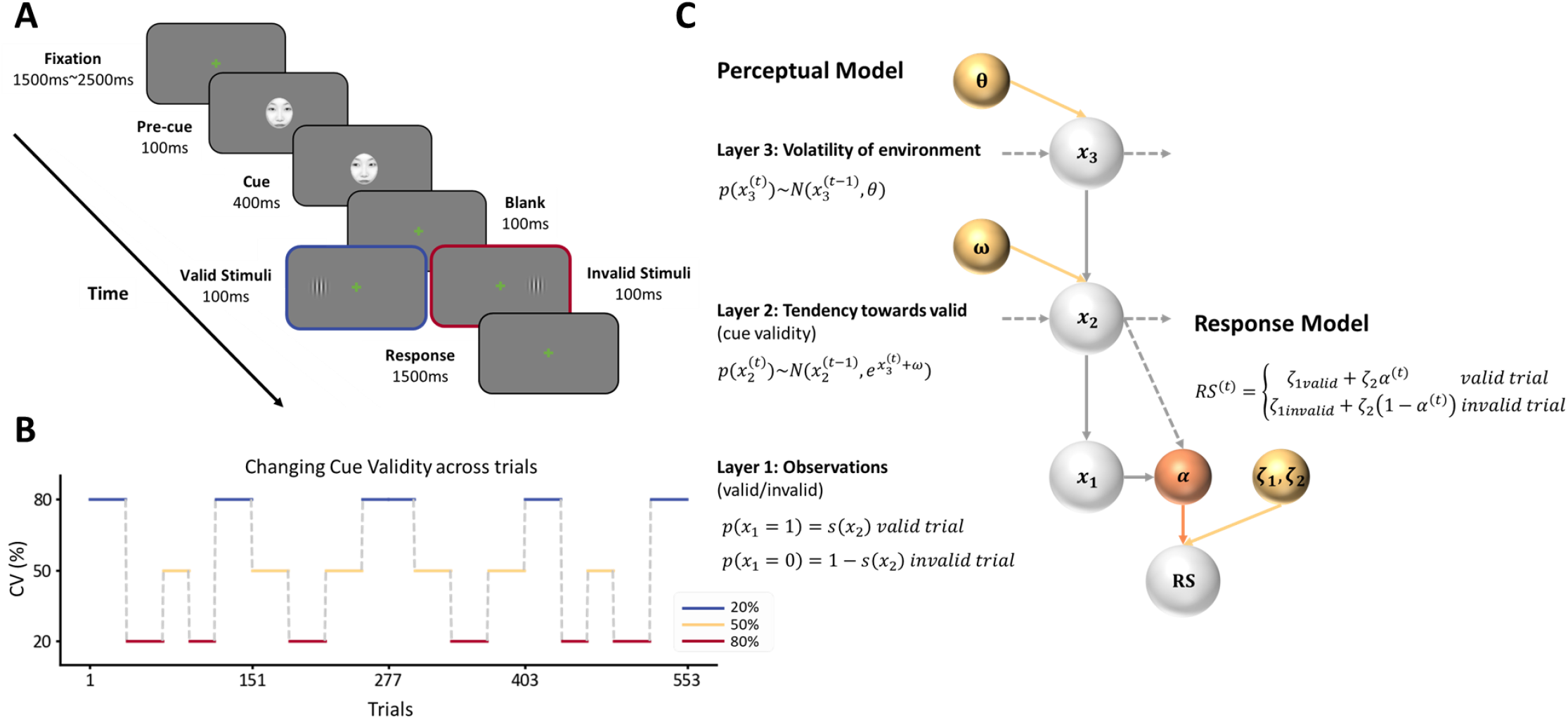
Experimental design and HGF model. (A) Modified Posner’s location-cueing paradigm. The social cue consisted of a human face, with eyes gazing left or right, predicting the position of the target grating. An example face image (for demonstration purposes only) is shown. Photo credit: Wenhui Gao, Beijing Normal University. Valid cues (blue box) matched the target position, while invalid cues (red box) did not. (B) CV Changed across trials. The solid-colored lines represented highly valid (blue line, CV = 0.8), highly invalid (red line, CV = 0.2), and noninformative (yellow line, CV = 0.5) cues, separately. (C) The HGF model comprises a perceptual model and a response model. The perceptual model includes three hidden layers (*x*_1_, *x*_2_, and *x*^3^), with belief updating influenced by subject-specific constants (ω and *θ*). The response model determines RS based on the precision-dependent attentional factor αand subject-specific constants (ζ_1_ and ζ_2_).

Each session consisted of 9 blocks, with a one-minute rest included after five blocks. Each block included 20 or 30 trials, with 4 catch trials randomly intermixed to maintain participants’ attention, resulting in a total of 552 trials. CV levels (20%, 50%, or 80%) varied across blocks (Fig. 1B). The gaze directions of the social cues were counterbalanced across blocks.

Participants were informed that CV levels would change during the experiment but were not given specific details about the timing or levels of these changes. The trial sequence was identical for all participants, a standard arrangement in computational studies of trial-wise learning (Behrens et al., 2007; Dyjas et al., 2012). This standardization ensures that variations in model parameters were due to individual differences rather than task-specific factors.

Before the main experimental sessions, participants completed practice blocks of 24 trials each, with a 50% CV, to ensure task familiarity. All participants achieved an accuracy above 97% and a median reaction time (RT) below 330 ms within 1–3 blocks.

### Behavioral analysis and modeling

#### Classical inference

Incorrect responses, misses, anticipations (RT < 150 ms), and RTs beyond 2 standard deviations (SDs) from each participant’s mean RT were excluded from the classical analysis and treated as null data in the Bayesian modeling. To meet the requirement of Bayesian model estimation, which assumes normally distributed variables, RTs were converted into response speed (RS, defined as the reciprocal of RT) to follow a normal distribution (Carpenter & Williams, 1995; Brodersen et al., 2008).

To investigate the impact of cue and its probabilistic context (CV) on RS, a 2 × 3 within-subject ANOVA was conducted with factors of cue (valid, invalid) and CV (20%, 50%, 80%). A significant cue × CV interaction indicated that probabilistic context influenced the cueing effect (i.e., RS_valid_ – RS_invalid_). Simple effects analysis with Bonferroni correction was performed to assess the significance of the cueing effect at each CV level. Furthermore, paired t-tests with Bonferroni correction were used to compare the cueing effects across different CV conditions.

#### Computational modeling

Bayesian models have been proved to offer a robust framework for explaining how individuals adjust their behavior based on probabilistic contexts in volatile environments (Dayan et al., 1995; Friston, 2009; Mathys et al., 2011). A comprehensive Bayesian model typically comprises a perceptual model and a response model. Here we employed the Hierarchical Gaussian Filter (HGF) (open source code: https://www.tnu.ethz.ch/de/software/tapas; see Fig. 1C) as our perceptual model, which maps belief trajectories of hidden states to sensory inputs on a trial-by-trial basis under the Bayesian framework (Mathys et al., 2011). Complementing this, our response model describes how these beliefs are translated into observable behavior (RS data).

In the perceptual model, three hierarchically coupled hidden states *x*_1_, *x*_2_, and *x*_3_ represented the hierarchical structure of the environment. The states *x*_2_ and *x*_3_ evolve over time according to Gaussian random walks *N*(·), with higher-level states modulating the variance of the lower-level states:

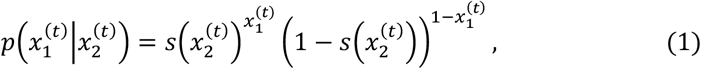

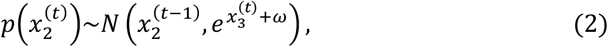

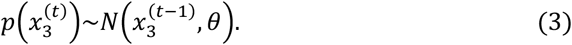

At the lowest level, 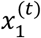 represents the observations from trial *t*, with possible outcomes 0 (invalid) or 1 (valid). The probability of 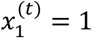 follows a Bernoulli distribution governed by the middle-level state 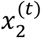 through a sigmoidal transformation *s*(·) (Eq. 1). The 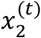 represents the inferred CV for trial *t*, which is normally distributed around 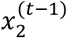. The variance of this distribution depends on both the state-dependent environmental volatility 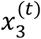 at the highest level and a parameter ω allowing for individual differences in updating 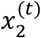 (Eq. 2). The state 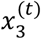 evolves following a normal distribution centered around its previous value 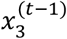, with variance determined by another subject-specific parameter *θ* (indicating the variability of the environmental volatility over time) (Eq. 3).

Following the principles of variational Bayes approximation (Beal, 2003; Mathys et al., 2011), participants’ trial-by-trial posterior belief updates 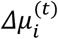 (i.e., 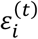) about each hidden state can be estimated (Daunizeau et al., 2010). The belief update is a function of the precision-weighted PEs at the lower level:

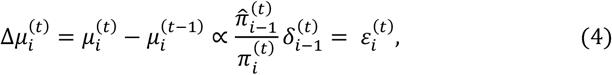

where 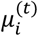 stands for the posterior means of the beliefs about the hidden states at level *i* for trial *t*. The hat symbol (*^*) indicates predictions prior to current observations. The 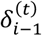 denotes the PEs at the lower level. The precision-weighted PEs are driven by the ratio of the prediction precision about the state at the lower level 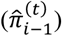 to the precision of the posterior belief at the current level 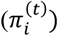. Apart from the first layer, the prediction precision at each layer is defined as the inverse variance of the prediction distribution. For the first layer, the prediction precision is defined based on CV predictions:

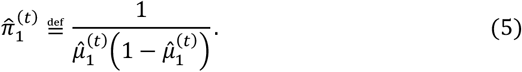

In the response model, participants’ posterior beliefs are mapped to the observed behavior (RSs). Here we used the “Precision” model as the response model, as detailed by (Vossel et al., 2014a). This model defines trial-wise RS as a linear function of the precision-dependent attentional factor α, which is determined by participants’ prediction precision at the first level of the perceptual model 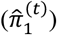. The equations are shown below:

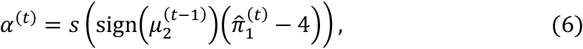

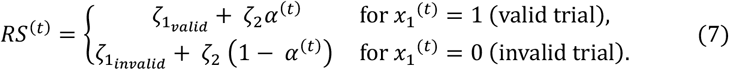

Here, the precision-dependent attentional factor αquantifies the allocation of attentional resources to the cued location and is derived via a sigmoid transformation (*s*(·)) of 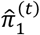, whose minimum value is 4 when the predicted CV 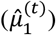 equals 0.5. The sign function (sign(·)) adjusts for counter-informative cues (when the 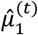 is below 0.5), ensuring that αranges between 0 and 1. 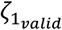 and 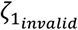 are subject-specific parameters, indicating the intercepts of RS for valid and invalid trials. The difference between 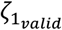 and 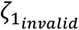 reflects the remaining exogenous cueing effects of social cues. ζ_2_ measures the influence of αon RSs, capturing the endogenous cueing effects of social cues.

#### Model comparison

To determine the optimal perceptual model for the observed data, we compared the specified three-level HGF (HSF_3_) against several alternative models. One alternative was a reduced version of HGF (HSF_2_), where the updating of the inferred CV was decoupled from the top level’s influence, similar to setting 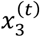 to zero.

To validate the “Precision” hypothesis on response mechanisms, we designed two alternative response models with distinct hypotheses about α(Vossel et al., 2014a). The “Belief” model posits that αis dependent on the prediction of CV (Eq. 8), whereas the “Surprise” model proposes that αis dependent on the Shannon surprise associated with the prediction of CV (Eq. 9):

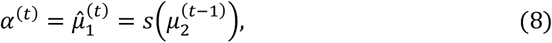

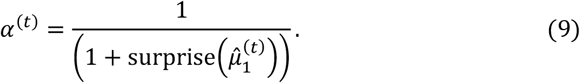

In addition, we included the Rescorla-Wagner (RW) model, a classical reinforcement learning framework (Rescorla & Wagner, 1972; see Eq. 10), which assumes that belief updating is driven by a fixed learning rate (α) and the trial-wise discrepancy between the observed reward (*R*) and the aggregated belief (∑ *V*):

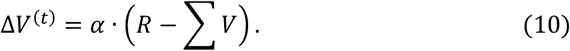

This RW model was directly integrated with the RS formulation (Eq. 7) by substituting α^*(t*)^ with the prediction of CV (*V*^*(t*)^).

These models formed a factorial model space comprising two hierarchical perceptual models and three response models, alongside an additional RW model. To evaluate the relative plausibility of these models at the group level, we employed random-effects Bayesian Model Selection (BMS) to compare the log-model evidences (LMEs) across models (Stephan et al., 2009). Specifically, the data from each participant were fitted to all the models, resulting in an LME for each model per participant. The exceedance probability (EP) and posterior probability (PP) from the BMS analysis were used, with EP assessing the relative likelihood that a given model is more frequent than all other models considered, and PP indicating the likelihood of each model given the data. The prior settings for all free parameters were detailed in Table 1.

**Table 1.** Prior and posterior distributions (mean and SD) of parameters in alternative models.

| Model |  | Perceptual model parameters |  | Response model parameters |  |  |
| --- | --- | --- | --- | --- | --- | --- |
| | | $\omega$ | $\theta$ | $\zeta_{1\text{valid}}$ | $\zeta_{1\text{invalid}}$ | $\zeta_2$ |
| HGF<sub>3</sub><br>"Precision" | Prior mean | -3.50 | -7.00 | 5.20E-03 | 5.20E-03 | 6.00E-05 |
|  | Prior SD | 0.50 | 0.50 | 0.1 | 0.1 | 1.00E-04 |
|  | Posterior mean | -3.47 | -7.00 | 3.79E-03 | 3.72E-03 | 6.00E-05 |
|  | Posterior SD | 0.12 | 2.97E-04 | 3.47E-04 | 3.44E-04 | 4.69E-09 |
| HGF<sub>3</sub><br>"Belief" | Prior mean | -3.50 | -7.00 | 5.20E-03 | 5.20E-03 | 6.00E-05 |
|  | Prior SD | 0.50 | 0.50 | 0.1 | 0.1 | 1.00E-04 |
|  | Posterior mean | -3.47 | -7.00 | 3.79E-03 | 3.72E-03 | 6.00E-05 |
|  | Posterior SD | 0.07 | 2.48E-04 | 3.47E-04 | 3.44E-04 | 4.16E-09 |
| HGF<sub>3</sub><br>"Surprise" | Prior mean | -3.50 | -7.00 | 5.20E-03 | 5.20E-03 | 6.00E-05 |
|  | Prior SD | 0.50 | 0.50 | 0.1 | 0.1 | 1.00E-04 |
|  | Posterior mean | -3.47 | -7.00 | 3.79E-03 | 3.72E-03 | 6.00E-05 |
|  | Posterior SD | 0.06 | 2.00E-04 | 3.47E-04 | 3.43E-04 | 3.05E-09 |
| HGF<sub>2</sub><br>"Precision" | Prior mean | -3.50 | -7.00 | 5.20E-03 | 5.20E-03 | 6.00E-05 |
|  | Prior SD | 0.50 | 0.50 | 0.1 | 0.1 | 1.00E-04 |
|  | Posterior mean | -3.44 | -7.00 | 3.79E-03 | 3.72E-03 | 6.00E-05 |
|  | Posterior SD | 0.13 | 2.97E-04 | 3.47E-04 | 3.43E-04 | 3.82E-08 |
| HGF<sub>2</sub><br>"Belief" | Prior mean | -3.50 | -7.00 | 5.20E-03 | 5.20E-03 | 6.00E-05 |
|  | Prior SD | 0.50 | 0.50 | 0.1 | 0.1 | 1.00E-04 |
|  | Posterior mean | -3.47 | -7.00 | 3.79E-03 | 3.72E-03 | 6.00E-05 |
|  | Posterior SD | 0.08 | 2.54E-04 | 3.47E-04 | 3.43E-04 | 3.85E-08 |
| HGF<sub>2</sub><br>"Surprise" | Prior mean | -3.50 | -7.00 | 5.20E-03 | 5.20E-03 | 6.00E-05 |
|  | Prior SD | 0.50 | 0.50 | 0.1 | 0.1 | 1.00E-04 |
|  | Posterior mean | -3.47 | -7.00 | 3.79E-03 | 3.72E-03 | 6.00E-05 |
|  | Posterior SD | 0.06 | 2.03E-04 | 3.47E-04 | 3.43E-04 | 2.94E-08 |
| | | $\alpha$ | $\nu$ | | | |
| RW | Prior mean | 0.05 | 0.5 |  |  |  |
|  | Prior SD | 1 | 0 |  |  |  |
|  | Posterior mean | 0.06 | 0.5 |  |  |  |
|  | Posterior SD | 0.02 | 0 |  |  |  |

To further analogize the model fitting results in the superior model with actual behavioral data, a multiple regression analysis was conducted to investigate the effects of cue and αvalues, as well as their interaction, on the RSs:

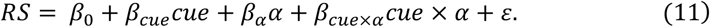

All individual data points were combined for a comprehensive analysis. The cue variable was dichotomously coded with -1 for invalid trials and 1 for valid trials. Both the continuous αand RS variables were standardized before analysis.

### fMRI data acquisition and analysis

#### MRI image acquisition and preprocessing

Structural and functional Magnetic Resonance Imaging (MRI) data were collected using a 3-Tesla Prisma MR Scanner (Siemens, Germany), equipped with a 64-channel head coil. High-resolution structural images were acquired using a T1 -weighted inversion recovery MPRAGE sequence (spatial resolution = 0.8 ×0.8 ×0.8 mm^3^, inversion time (TI) = 1000 ms, repetition time (TR) = 2400 ms). Functional images with BOLD contrasts were obtained using a whole-brain T2^*^-weighted echo-planar imaging (EPI) sequence (TR = 1500 ms, echo time (TE) = 30 ms, field of view (FoV) = 200 mm, spatial resolution = 2 ×2 ×2 mm^3^, 76 slices (axial, interleaved)). Each of the two functional runs during the main task session comprised 820 volumes (approximately 20 min 40 s of acquisition time).

To account for T1 equilibration effects, the initial five volumes from each functional run were discarded. Subsequent volumes were then processed using Statistical Parametric Mapping software (SPM12; www.fil.ion.ucl.ac.uk/spm, Wellcome Department of Imaging Neuroscience, London). Preprocessing included the following steps. First, functional images underwent temporal sinc interpolation for slice timing correction. The images were then realigned to the first volume using rigid body transformations to correct for head movements and generate 6 motion regressors and a mean functional image. Each participant’s structural image was coregistered to this mean functional image, followed by segmentation. The parameters obtained from segmentation were applied to normalize both structural and functional images to the Montreal Neurological Institute (MNI) template, with the functional images resampled to 2 ×2 ×2 mm^3^ voxel size. Finally, normalized functional images were spatially smoothed using a Gaussian kernel with a full-width at half maximum (FWHM) of 6 mm.

#### General linear model (GLM) analysis

To explore the neural correlates of attentional allocation, we performed model-based fMRI analyses. Specifically, we constructed three GLMs to independently explain each voxel’s BOLD signal time series with three parameter trajectories (α, |*ε*_2_|, |*ε*^3^|) derived from the winning computational model. These GLMs were constructed individually for each participant to account for the subject-specific nature of the parameter trajectories. The three GLMs are as follows:

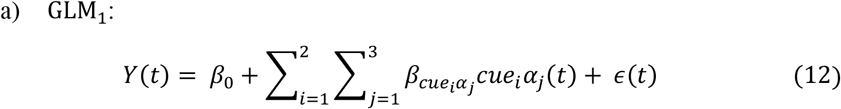

In GLM_1_ (Eq. 12), the factors of cue (valid, invalid) and categorized αlevels (low, medium, high) were used as predictors of BOLD responses (Y) at the first (individual) level. The αcategories were determined by dividing individual αvalues into tertiles. This analysis generated a contrast map for each cue and αlevel for each participant. These contrast maps were subsequently entered into a second (group) level within-subject random-effects ANOVA with a full factorial design. This approach enabled the detection of interaction effects between cue and the attentional factor α, identifying brain regions showing differential responses to invalid versus valid trials at different αlevels. Additionally, visualization of this whole-brain ANOVA analysis was performed at the region of interest (ROI) level. ROIs were defined using spheres with a 4 mm radius, centered on the peak coordinates of clusters demonstrating significant interaction effects. The percentage of signal change in these ROIs was estimated using MarsBaR-0.42 (Brett et al., 2002) for a detailed interpretation of the observed interaction effects.

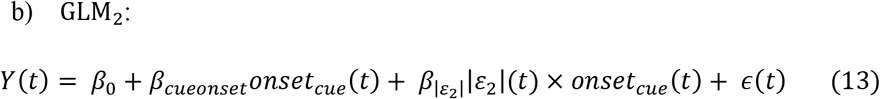

In GLM_2_ (Eq. 13), the trajectory of the absolute precision-weighted PEs about target locations (|*ε*_2_|) were incorporated as a computational parametric modulator. This model aimed to identify brain regions where the BOLD response amplitude varied as a function of the modulation of |*ε*_2_| at cue onset. The absolute value of ε_2_ was chosen because the magnitude of the PEs is more closely related to the learning process (Fouragnan et al., 2024) and can elicit stronger responses in the attentional network (Fouragnan et al., 2017) compared to the signed PEs. At the group level, t-contrasts were employed to test the significance of the modulation effect (parametric modulation effect > baseline).

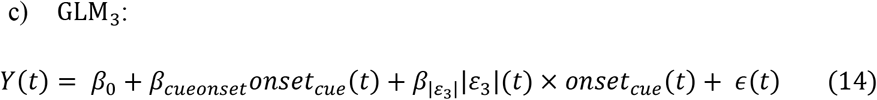

Similar to GLM_2_ (Eq. 13), GLM_3_ (Eq. 14) used the trajectories of the absolute precision-weighted PEs about CV (|*ε*^3^|) as computational parametric modulator. This model focused on detecting brain regions sensitive to the modulation of |*ε*^3^| at cue onset.

Additionally, another GLM was constructed to examine the main effect of cue (valid, invalid) on BOLD responses.

All GLMs included catch trials and trials discarded from behavioral analysis as separate regressors. Confounding regressors, including temporal derivatives and a comprehensive set of 24 head motion parameters, were also incorporated into the GLMs. The 24 motion parameters consisted of the 6 standard motion parameters, their first temporal derivatives, 6 squared standard parameters, and the temporal derivatives of these squares (Friston et al., 1996; Yan et al., 2013). All regressors were convolved with a canonical hemodynamic response function (HRF).

Statistical significance for all reported results was determined using a threshold of *p* < .05 with cluster-level FDR correction after a voxel-level cutoff of *p* < .0005 (uncorrected).

#### Dynamic causal modeling (DCM)

We further employed DCM (Friston et al., 2003; Stephan et al., 2010) to examine the modulation of |*ε*_2_| on the strength of the connections between brain regions where BOLD responses to cue onsets were significantly modulated by |*ε*_2_|. Specifically, we focused on the primary visual cortex (V1), STS, and TPJ. Time series data from these regions were extracted for each participant across two experimental runs and concatenated for DCM analysis.

In the context of neuronal responses to visual inputs, V1 has been consistently identified as the primary recipient of visual stimuli, as supported by both electrophysiological findings (Felleman & Van Essen, 1991) and DCM analysis (Raffin et al., 2022). Additionally, studies have demonstrated that the STS (Sadeghi et al., 2022) and TPJ (Vossel et al., 2015) are also capable of receiving visual inputs under certain conditions. Based on this evidence, we specified three model families positing driving inputs to: (1) V1 and STS; (2) V1 and TPJ; or (3) V1, STS, and TPJ. Each model family included a “full” DCM model with reciprocal connections among the regions and intrinsic self-connections, all modulated by |*ε*_2_|.

The optimal model explaining the observed responses in V1, STS, and TPJ was determined using Bayesian model reduction (Friston et al., 2016) and BMS. First, family-level inference (Penny et al., 2010) was applied to determine the most appropriate entry points for visual inputs. Next, the models within the winning family were compared to pinpoint the most plausible DCM that accounted for the modulation effects of |*ε*_2_|. Bayesian model averaging was then applied to integrate the parameters from all reduced models in the winning model family, weighted by their model evidence. Modulatory connectivity parameters were considered highly significant if their PP exceeded .99 and moderately significant if they exceeded .90.

## Results

### Behavioral results

Participants’ overall accuracy in the task was remarkably high, averaging 98.66% (±1.04% SEM), with 98.79% (±1.22% SEM) in the first run and 98.53% (±1.68% SEM) in the second. After excluding trials with incorrect responses, misses, anticipations, and deviating RTs, 94.96% (±1.56% SEM) of the trials across participants were retained for analysis.

The 2 (cue: valid, invalid) ×3 (CV: 20%, 50%, 80%) within-subject ANOVA on mean RSs revealed a significant interaction effect (*F*_(2.64)_ = 9.45, *p* < .001,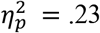), indicating that the differences in RSs between valid and invalid trials varied across different CV levels (Fig. 2A). Further simple effects analysis showed significant differences between valid and invalid trials at each CV level: 20% (*F*_(1,32)_ = 7.43, *p* < .01,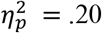), 50% (*F(*1,_3_2) = 24.94, *p* < .001,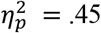), and 80% (*F(*1,_3_2) = 56.34, *p* < .001,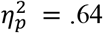). Further paired t-tests were conducted to directly examine the impact of CV on the cueing effect (RS_valid_ – RS_invalid_). The analyses revealed significant differences between CV levels of 20% versus 80% (*t*_(32)_ = 4.50, *p* < .001, Cohen’s *d* = .79), and 50% versus 80% (*t*_(32)_ = 2.89, *p* < .05, Cohen’s *d* = .52), but not of 20% versus 50% (*t*_(32)_ = 1.60, *p* = .118 > .05, Cohen’s *d* = .31). These results indicate that the cueing effect is significantly stronger at the higher CV level of 80% compared to the lower CV levels of 20% and 50%.

**Fig. 2.**
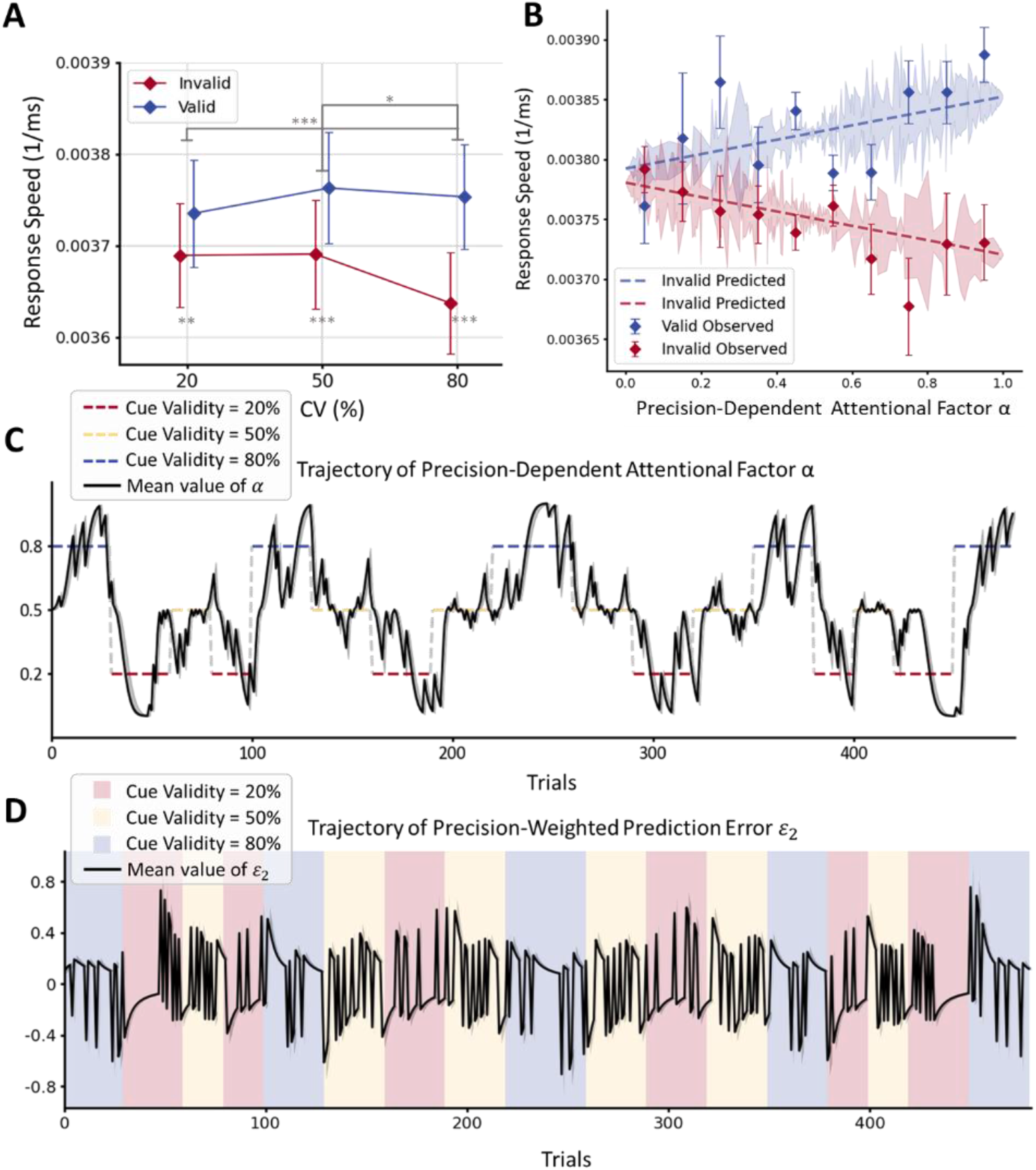
Behavioral results. (A) Mean RSs for each cue and CV condition. Asterisks below indicate significant differences between valid (blue diamond) and invalid (red diamond) trials. Asterisks above indicate significant differences in cueing effects between CV conditions. Multiple comparisons were adjusted by Bonferroni correction. (B) Observed and predicted RSs as a function of α. Observed RSs were clustered into ten bins based on αvalues, separately for valid (blue) and invalid (red) trials. Predicted RSs were computed based on Eq. 7. (C) Trajectory for group average α(solid black line). Colored dashed line represents time-varying CV. (D) Trajectory for group average ε_2_ (solid black line). Colored shadow shows time-varying CV. All the parameters were estimated from the HSF_1_ “Precision” model. Error bars and colored shades represent standard errors. ^*^ .01 < *p* ⩽ .05, ^**^ .001 < *p* ⩽ .01, ^***^ *p* ⩽ .001.

### Behavioral modeling

We employed a Bayesian model to characterize the influence of the volatile probabilistic cue contexts on individual behavioral responses. Specifically, we used the HGF (Mathys et al., 2011) as the perceptual model, paired with a response model. We compared two versions of the HGF: HSF^3^, where beliefs about CV volatility (*x*^3^) are updated trial-by-trial and affect the update of CV beliefs (*x*_2_), and HSF_2_, where CV volatility does not affect CV belief updates. Within the response model framework, we compared three hypotheses: the “Precision” model, the “Belief” model, and the “Surprise” model. In addition, we included a standard reinforcement learning model— RW model for comparison with the Bayesian models. All estimated posterior values of the free parameters for each model were presented in Table 1.

A random-effects BMS (Table 2) indicated that the HSF_3_ model family outperformed the HSF_2_ family among the perceptual models (EP > .99, PP = .99), and the “Precision” model family was the winning model family among the response model families (EP > .99, PP = .98). Finally, a comprehensive comparison of all 7 individual models identified the combined HSF_3_ and “Precision” model as superior to all other models (EP > .99, PP = .98).

**Table 2.** Results of Bayesian model selection (BMS)

| Model | PP | EP |
| --- | --- | --- |
| Model family comparison: perceptual models |  |  |
| HGF<sub>3</sub> family | 0.985 | 0.999 |
| HGF<sub>2</sub> family | 0.015 | < 0.001 |
| Model family comparison: response models |  |  |
| “Precision” family | 0.980 | 0.999 |
| “Belief” family | 0.010 | < 0.001 |
| “Surprise” family | 0.010 | < 0.001 |
| Model comparison of all 7 models |  |  |
| HGF<sub>3</sub> “Precision” | 0.975 | 0.999 |
| HGF<sub>3</sub> “Belief” | 0.004 | < 0.001 |
| HGF<sub>3</sub> “Surprise” | 0.004 | < 0.001 |
| HGF<sub>2</sub> “Precision” | 0.004 | < 0.001 |
| HGF<sub>2</sub> “Belief” | 0.004 | < 0.001 |
| HGF<sub>2</sub> “Surprise” | 0.004 | < 0.001 |
| RW | 0.004 | < 0.001 |

PP, posterior probability; EP: exceedance probability.

We plotted participants’ average trial-wise estimates of precision-dependent attentional factor α(Fig. 2C; Eq. 6) and precision-weighted PEs about target locations (ε_2_) (Fig. 2D; Eq. 4) from the winning model. The trajectory of αclosely followed the fluctuations in CV, indicating that attentional allocation to the cued location varied consistently with CV. Moreover, the estimated RSs as a function of binned αand cue were visualized (Fig. 2B; Eq. 7). A multiple regression analysis revealed a significant interaction between cue and α(β_*cue*×α_ = .036, *p* < .001) and a main effect of cue (β_*cue*_ = .056, *p* < .001), confirming the alignment between the behavioral data and the model estimates. Consistent with the model comparison results, the estimated RSs from the winning model provided the best fit to the observed RS data, as evidenced by the highest LME (Fig. S1).

### Neural correlates of precision-dependent attentional factor α

To investigate the neural correlates of precision-dependent attentional allocation, a voxel-wise whole-brain ANOVA of activation based on the cue (valid, invalid) and categorized α levels (low, medium, high) interaction was performed as illustrated in GLM_1_. This analysis revealed significant interaction effects in the bilateral MTG, right superior frontal gyrus (SFG), left insula, left IPL, left MFG, and right IFG (Fig. 3; Table 3). Notably, pronounced interaction effects were observed in the right TPJ within the supramarginal gyrus, and the right FEF within the PreCG. Moreover, significant interaction effects were also associated with activity in regions including the right fusiform face areas (FFA), STS, and IPS. Importantly, no region exhibited a significant main effect for either cue or categorized α levels. To directly compare with traditional findings in spatial attention research (Kuhns et al., 2017; Vossel et al., 2015), we conducted a voxel-wise one-way ANOVA to assess the main effect of cue (valid, invalid) on whole-brain activity. No significant regions were identified, likely due to the reversal of neural activity when CV dropped below 50%.

**Table 3.** Neural correlates of the interactions between cue and precision-dependent attentional factor α.

| Region | Hemisphere | MNI coordinates |  |  | Voxels | *F*-score |
| --- | --- | --- | --- | --- | --- | --- |
|  |  | x | y | z |  |  |
| MTG / STS | R | 42 | -64 | 10 | 1362 | 18.91 |
| MTG | L | -48 | -66 | 8 | 452 | 17.71 |
| SFG | R | 6 | 8 | 54 | 508 | 17.63 |
| Insula | L | -46 | -2 | 12 | 458 | 16.43 |
| TPJ | R | 56 | -34 | 30 | 278 | 15.78 |
| FFA | R | 38 | -56 | -18 | 370 | 15.75 |
| IPL | L | -54 | -26 | 24 | 553 | 14.22 |
| MFG | L | -32 | -6 | 66 | 82 | 14.13 |
| IPS | R | 22 | -54 | 44 | 112 | 13.10 |
| IFG | R | 54 | 16 | 0 | 99 | 12.61 |
| FEF | R | 44 | -4 | 54 | 174 | 12.54 |

All results were significant with $p < .05$ FDR cluster-level corrected. R, right; L, left; MTG, medial temporal gyrus; STS, superior temporal sulcus; SFG, superior frontal gyrus; TPJ, temporo-parietal junction; FFA, fusiform face area; IPL, inferior parietal lobule; MFG, middle frontal gyrus; IPS, intraparietal sulcus; IFG, inferior frontal gyrus; FEF, frontal eye fields.

**Fig. 3.**
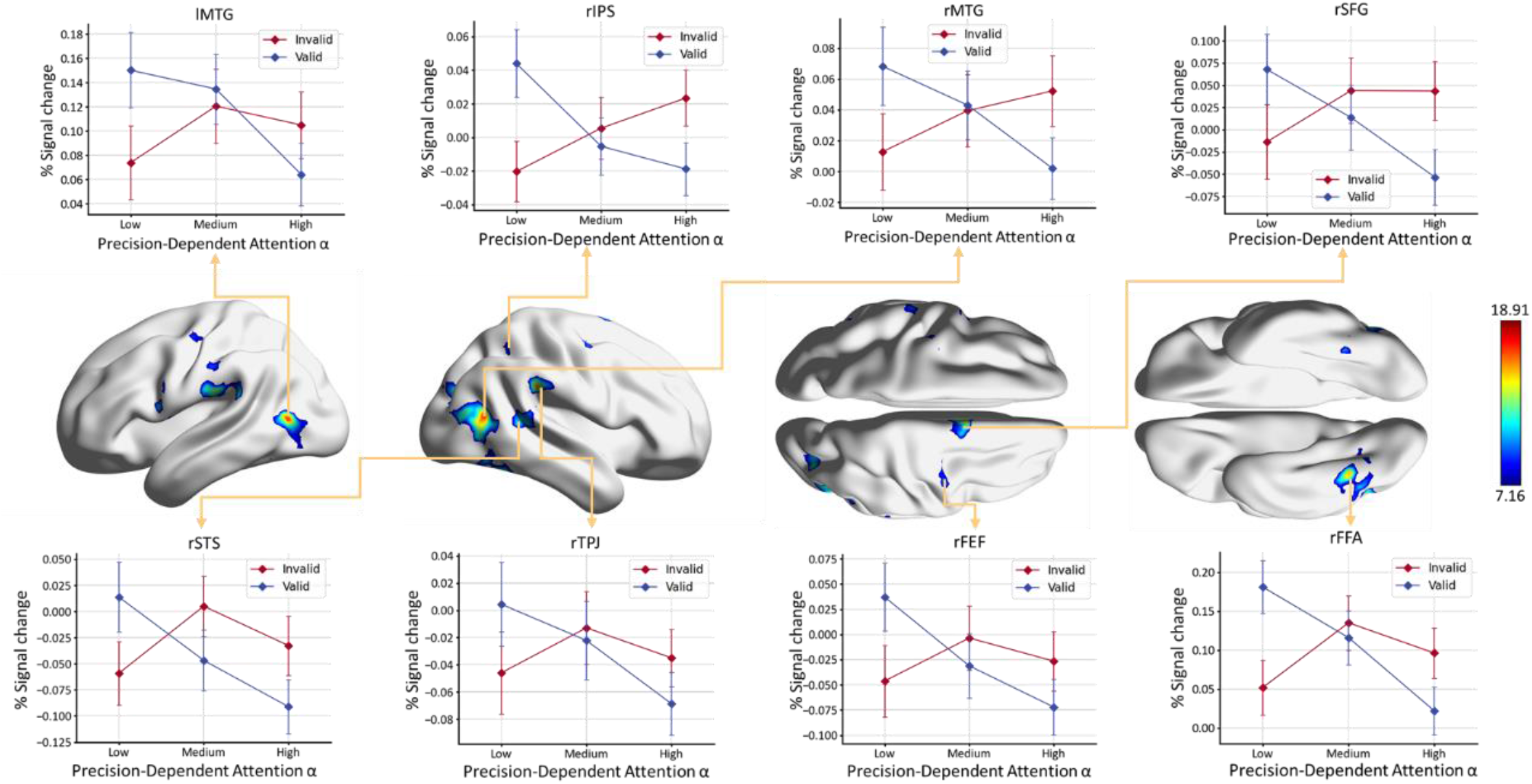
Whole-brain activation for interaction effects between CV and αlevels (p < .05, cluster-level FDR corrected). Middle panel: Significant interactions between αlevels (low, medium, and high) and cue (valid, invalid). The color bar represents the *F*-statistic range. Top and bottom panels: Mean percentage change in BOLD signal within the ROIs under valid (blue diamond) and invalid (red diamond) conditions. Error bars represent standard errors. l, left hemisphere; r, right hemisphere. MTG, middle temporal gyrus; IPS, intraparietal sulcus; SFG, superior frontal gyrus; STS, superior temporal sulcus; TPJ, temporo-parietal junction; FEF, frontal eye fields; FFA, fusiform face areas.

To further quantify the cue ×αinteraction effects, percentages of signal changes at the ROI level were extracted for each participant (Fig. 3). These ROIs were defined centered on peak coordinates from regions exhibiting significant interaction effects (Table 3). Nearly all ROIs exhibited an X-shaped activation pattern, indicating a reversal of the cueing effect on activations as αincreased from low to high levels. This pattern suggested that the influence of cue on BOLD activities is modulated by the precision-dependent attention.

### Modulation effects of precision-weighted PEs on brain activity and inter-regional connectivity

In a volatile environment characterized by changes in CV, the efficient allocation of attention was guided by updated inferences on environmental statistics. According to the HGF model, these updates were primarily driven by ε_2_ and ε_3_ (Mathys et al., 2011). To effectively identify brain regions involved in this process, parametric modulation was employed at the whole-brain level using the absolute values of these precision-weighted PEs (|*ε*_2_| in GLM_2_ and |*ε*^3^| in GLM^3^) as modulators. This analysis demonstrated significant BOLD response increases correlated with higher |*ε*_2_| in V1, STS, and TPJ (Fig. 4A; Table 4). The modulation effects in these regions highlighted their pivotal roles in adjusting inferences in response to CV changes, facilitating efficient attentional allocation. However, no any region was significantly modulated by |*ε*^3^|.

**Table 4.** Neural correlates of the modulation effect of absolute precision-weighted PE about CV (|*ε*_2_|)

| Region | Hemisphere | MNI coordinates |  |  | Voxels | *t*-score |
| --- | --- | --- | --- | --- | --- | --- |
|  |  | x | y | z |  |  |
| V1 | R | 4 | -70 | 2 | 574 | 5.75 |
| V1 | L | -26 | -96 | -2 | 72 | 5.21 |
| STS | R | 32 | -68 | 24 | 46 | 4.71 |
| TPJ | R | 60 | -36 | 28 | 66 | 4.34 |

All results were significant with $p < .05$ FDR cluster-level corrected. R, right; L, left; V1, primary visual cortex; STS, superior temporal sulcus; TPJ, temporo-parietal junction.

**Fig. 4.**
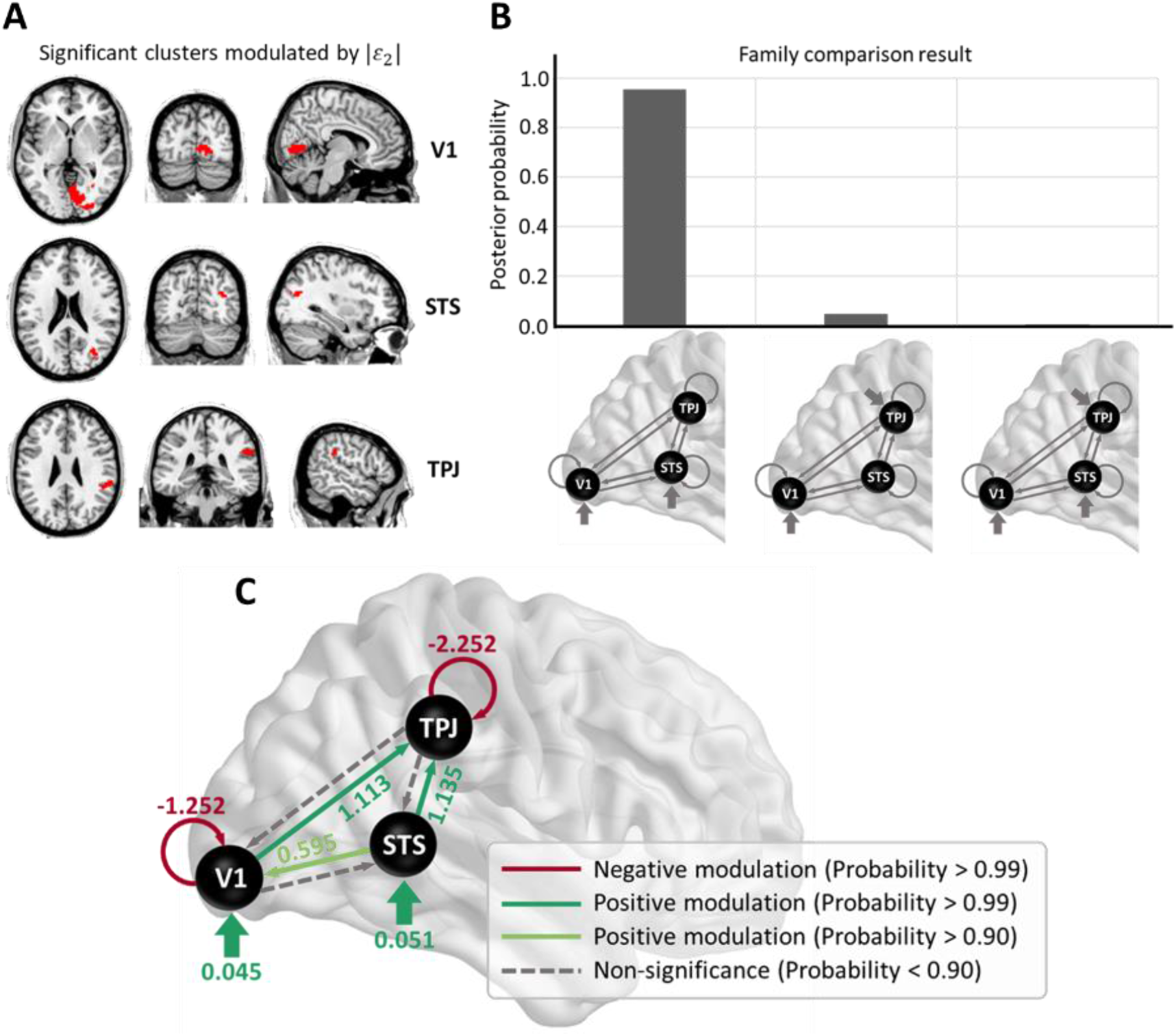
DCM results. (A) Neural correlates of the modulation effect of |*ε*_2_| (*p* < .05, cluster-level FDR corrected). The three regions significantly modulated by |*ε*_2_| including V1, STS, AND TPJ, were entered in the DCM analysis. (B) Results of the DCM family comparison. The posterior probabilities for the three model families after Bayesian model reduction (gray bars). The winning model family, with the highest posterior probability, posited driving inputs to both V1 and STS. (C) Mean modulatory parameter estimates for |*ε*_2_|. Positive driving inputs (bold green arrows) were directed to V1 and STS. Solid arrows indicate connections with modulatory effects (red: negative, posterior probability > .99; green: positive, posterior probability > .99; light green: positive, posterior probability > .90). V1, primary visual cortex; STS, superior temporal sulcus; TPJ, temporo-parietal junction.

To further elucidate the directionality and the extent to which |*ε*_2_| modulated the effective connectivity among the identified regions, DCM (Friston et al., 2003; Stephan et al., 2010) was applied to BOLD responses. An initial family comparison revealed that the superior configuration involved inputs to V1 and STS, with a PP equals .96 (Fig. 4B). Subsequent Bayesian model reduction within the winning family identified the best-fitting model and enabled a detailed evaluation of its parameters (Fig. 4C). The results indicated that |*ε*_2_| significantly enhanced the influence of STS on both V1 and TPJ, as well as the impact of V1 on TPJ. Concurrently, |*ε*_2_| diminished the self-influences of V1 and TPJ. Notably, all parameters exhibited PPs exceeding .90.

## Discussion

The current study employed a hierarchical Bayesian model to investigate the computational and neural mechanisms underlying context effects in social attention. Behavioral findings demonstrated that the gaze cueing effect increased with cue informativeness and remained significant even when the cues were counter-informative. Bayesian modeling analysis showed that individuals generated hierarchical predictions based on dynamic environmental statistics, with social attention modulated by the precision of predictions regarding CV. fMRI results further revealed that brain regions involved in spatial attention and social perception were associated with precision-dependent attentional allocation. Precision-weighted PEs about target locations influenced activity in V1, STS and TPJ, as well as their effective connectivity, underscoring their roles in updating predictions.

This study extended previous spatial attention paradigms with a broad range of CV levels (Vossel et al., 2014a, 2014b, 2015; Kuhns et al., 2017; Li et al., 2023), adapting them to social attention. The context effect of CV on gaze cueing observed here aligned with earlier findings from arrow cueing studies (Vossel et al., 2014a, 2014b). Notably, the cueing effect remained significant even at a counter-informative 20% CV level, consistent with previous research on social cues (Friesen et al., 2004; Tipples, 2008). Two potential explanations for the persisting cueing effect at CV levels below 50% were considered: (1) a biased belief that overestimated the actual CV; or (2) an unbiased belief where social cues reflexively influenced attention shifts. Bayesian model fitting analysis indicated that individuals’ predictions of CV closely matched the actual CV (Fig. S2), ruling out the first explanation. This suggested that the observed cueing effect was more likely driven by the reflexive component of social attention (Friesen & Kingstone, 1998; Driver et al., 1999; Friesen et al., 2004). Consequently, our findings implied that behavioral responses to social cues reflected not only theoretically accurate voluntary attentional allocation but also potential reflexive aspects of attention.

The hierarchical Bayesian model played a crucial role in characterizing the computational mechanisms underlying voluntary social attentional allocation. This study demonstrated that attentional allocation was modulated by the precision of predictions for CV, aligning with previous research on spatial attention (Feldman & Friston, 2010; Vossel et al., 2014a, 2014b, 2015). Within the Bayesian framework, precision referred to the certainty in predictions about environmental states (Feldman & Friston, 2010), which guided attentional allocation (Rao, 2005). As prediction precision increased, individuals relied less on immediate sensory evidence and more on prior information (Yu & Dayan, 2005; Kiani et al., 2014).The winning model (HSF^3^) (Fig. S3) showed that when cue predictability 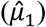 was near 0.5, the prediction precision of CV 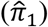 was low, resulting in a more evenly distributed allocation of attention (αnearing 0.5). Conversely, when cue was highly predictive (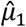 approaching 0 or 1), prediction precision 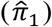 increased, concentrating attention on the predicted target location. This implied that attention is more efficiently allocated to highly predictable social cues, optimizing the use of cognitive resources. The model further suggested that the brain extracts hierarchical environmental statistics from visual experiences (Vossel et al., 2014a; Diaconescu et al., 2017; Sevgi et al., 2020; Hein et al., 2021), consistent with sparse coding principles, which efficiently reduce the dimensionality of complex sensory inputs (Beyeler et al., 2019) and facilitate information processing from sensory areas to higher-level frontal regions (Elmoznino & Bonner, 2024). This hierarchical processing explained why the HGF model more accurately reflected individual performance than non-hierarchical models.

Further analysis identified key brain regions involved in voluntary social attentional allocation, particularly in response to the interaction between attention and CV. The results showed significant activation in regions traditionally associated with attention, including the TPJ, FEF, and IPS, along with face perception areas like the STS and FFA. The TPJ, a central node in the VAN (Corbetta et al., 2008), was crucial for reorienting attention towards task-relevant but previously unattended stimuli (Arrington et al., 2000; Corbetta & Shulman, 2002), a role supported by its involvement in prediction updating (Geng & Vossel, 2013) and validated through the transcranial magnetic stimulation (TMS) study (Mengotti et al., 2017). The IPS and FEF, components of the DAN (Sato et al., 2009), were involved in generating and transmitting spatial attention predictions through top-down signals to visual areas (Corbetta & Shulman, 2002; Monosov et al., 2008; Moore & Armstrong, 2003; Sato et al., 2016). Moreover, the STS and FFA, which were critical for processing dynamic and static facial features respectively (Hasselmo et al., 1989; Haxby et al., 2000), were traditionally considered key regions in social perception. The current study extended this by demonstrating their involvement in precision-dependent social attention. This finding aligned with previous research showing that both regions were significantly engaged in processing gaze cues (Nummenmaa et al., 2010; Nummenmaa & Calder, 2009), with the FFA primarily involved in gaze processing (Andrews & Ewbank, 2004) and the STS in decoding intentions conveyed through gaze (Hooker et al., 2003; Mosconi et al., 2005). Thus, both regions could be modulated by the direction of social attention. Taken together, these findings suggested that social attention is governed by the integration of spatial attention with social perception processing.

Trial-by-trial absolute precision-weighted PEs about target locations (|*ε*_2_|), reflecting the adjustment magnitude needed to align predictions with actual occurrence of target locations (Fouragnan et al., 2017; Rouhani & Niv, 2021), were favored over signed PEs, which indicated reinforcement or extinction of previous choices (Dayan & Daw, 2008). Our analysis showed significant associations between |*ε*_2_| and activations in regions including V1, TPJ and STS, underscoring their roles in updating environmental predictions. Previous research on non-social spatial attention has demonstrated that |*ε*_2_| modulated activities in the frontal gyri, ACC, IPS, anterior insula, and dopaminergic midbrain (Diaconescu et al., 2017; Iglesias et al., 2021), as well as gamma-band activity in V1 and TPJ (Auksztulewicz et al., 2017). The current evidence further highlighted the crucial role of the STS in processing social cue-related PEs, thus facilitating the updating of predictions.

DCM analysis provided deeper insights into how |*ε*_2_| influenced effective connectivity across these regions involved in social attention. Specifically, visual stimuli were predominantly conveyed to and processed in V1 and STS, consistent with prior studies (Felleman & Van Essen, 1991; Sadeghi et al., 2022). The increased |*ε*_2_| strengthened the forward connections from V1 and STS to TPJ (Friston et al., 2003; Stephan et al., 2010), suggesting that these connections might transmit residual errors, thereby enhancing the role of TPJ in processing prediction violations and updating predictions (Vossel et al., 2012, 2015). This process aligned with the predictive coding theory, where sensory regions generated PEs that were forwarded to the TPJ and other higher cortical areas for further processing and adjustment of predictions (Rao & Ballard, 1999). These findings emphasize the TPJ’s centrality in the dynamic interaction between sensory input and higher-order prediction mechanisms, highlighting its critical role in modulating attention based on the precision of sensory information.

Several questions remained for further investigation. The temporal resolution of fMRI might have been insufficient to differentiate cue-locked prediction modulation from target-locked PE modulation, potentially reducing effect sizes. Techniques like MEG, with higher temporal resolution, could more effectively distinguish these neural processes (Auksztulewicz et al., 2017). Moreover, the absence of direct comparisons between social and endogenous cues limited our ability to identify regions specifically involved in social attention—a gap that future research should address.

Understanding the neural mechanisms of social attention could inform interventions for conditions like ASD (Lawson et al., 2017), ADHD (Hauser et al., 2014), and schizophrenia (Stephan & Mathys, 2014; Diaconescu et al., 2019), where prediction updating is impaired. The application of the hierarchical Bayesian model to social attention could guide targeted treatments. Furthermore, insights from social attention research could enhance AI development, particularly in social intelligence (Fan et al., 2022). In conclusion, this study not only explored the computational and neural mechanisms of social attention but also provided a foundation for future research in clinical and AI domains.

## Code and data availability

All materials and data have been made publicly available via the Open Science Framework (https://osf.io/vraqp/).

## Acknowledgments

This work was supported by the STI2030-Major Projects (2021ZD0203803), the National Natural Science Foundation of China (32200840), National Key R&D Program of China (2019YFA0709503), China Postdoctoral Science Foundation (2022T150061, 2022M710435), and Fundamental Research Funds for the Central Universities.

## Author contributions

W.G. was responsible for data curation, formal analysis, investigation, visualization, and writing – original draft, and writing – review & editing. W.G. and L.Z. were responsible for writing – review & editing. W.G., L.Z. and K.Z. were responsible for conceptualization and validation. C.Z. and B.S. were responsible for methodology. K.Z. and L.Z. were responsible for funding acquisition. B.S., L.Z. and K.Z. were responsible for supervision.

