## Supplemental Figure 1 for "Precision-dependent modulation of social attention"

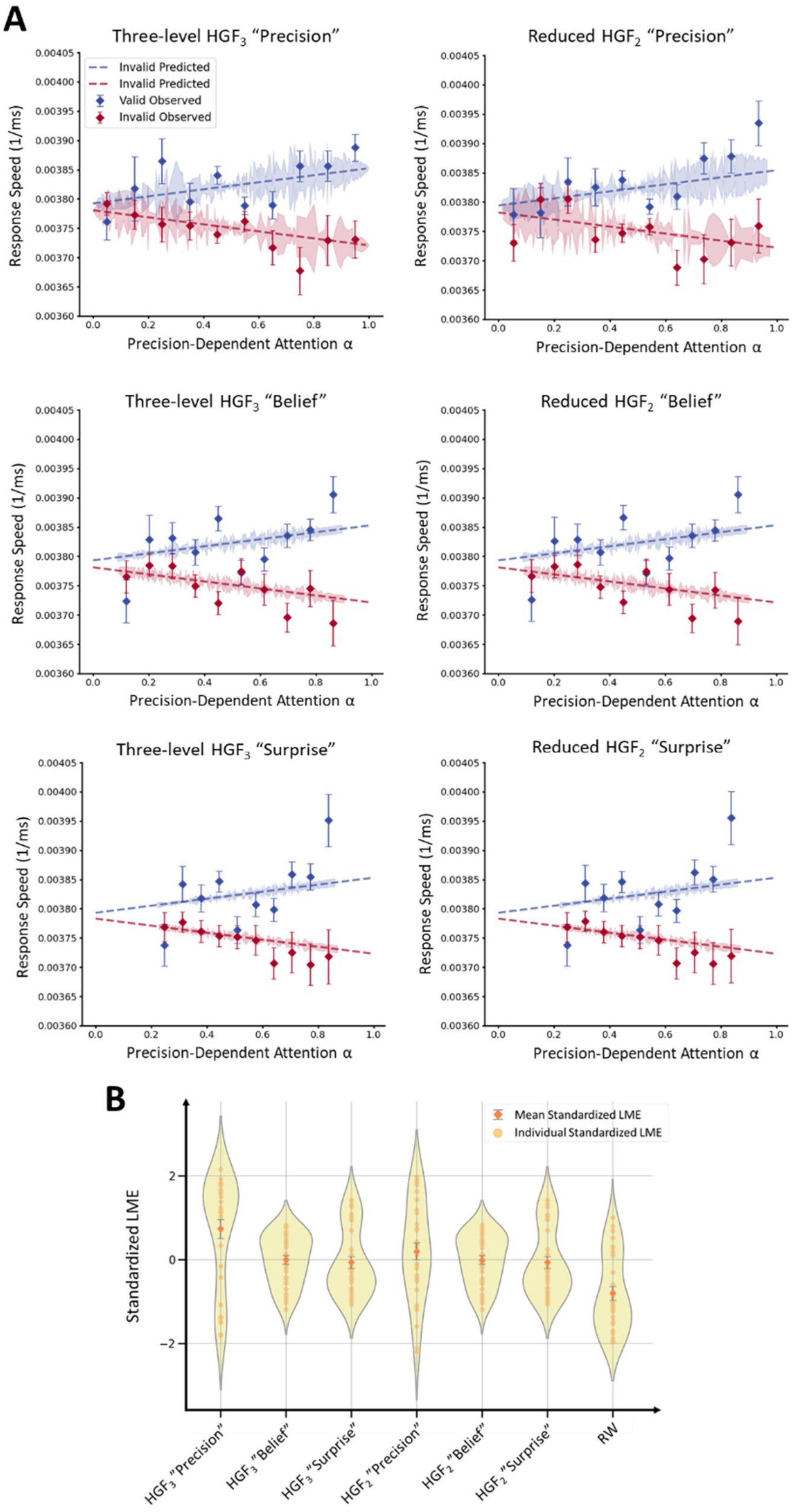


Fig. S1. (A) Observed and predicted RSs as a function of $\alpha$ across computational models. The top row presents the results from $HGF_3$ and the bottom row presents the results from $HGF_2$. Each column, from left to right, corresponds to the results from the "Precision" model, the "Belief" model, and the “Surprise” model. Observed RSs were clustered into ten bins based on $\alpha$ values, separately for valid (blue) and invalid (red) trials. Predicted RSs were computed based on Eq. 7. (B) Standardized LMEs across computational models. LMEs were computed by comparing model estimates with actual data and standardized for each individual across 7 alternative models (yellow dots). The mean standardized LMEs in each model are represented by orange diamonds. Error bars represent standard error.
