## Supplemental Figure 2 for "Precision-dependent modulation of social attention"

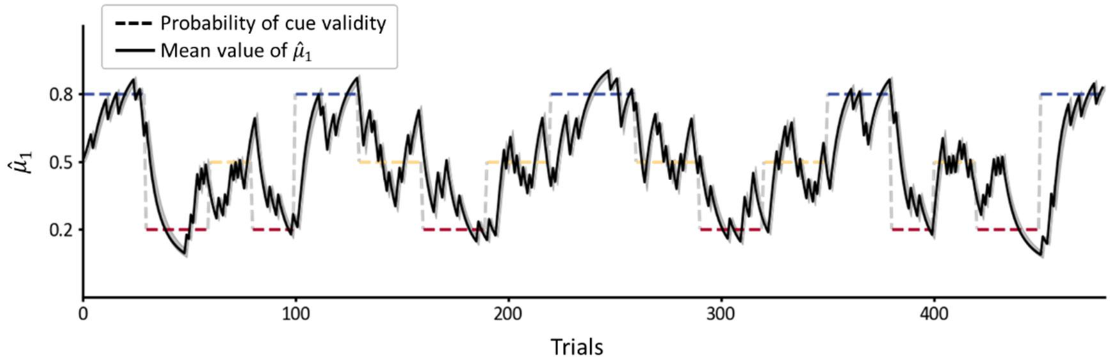


Fig. S2. Trajectory for group average $\hat{\mu}_1$ (solid black line). Colored dashed line represents time-varying CV. Parameters were estimated from the $HGF_1$ "Precision" model. Gray shades represent standard errors.
