## Supplemental Figure 3 for "Precision-dependent modulation of social attention"

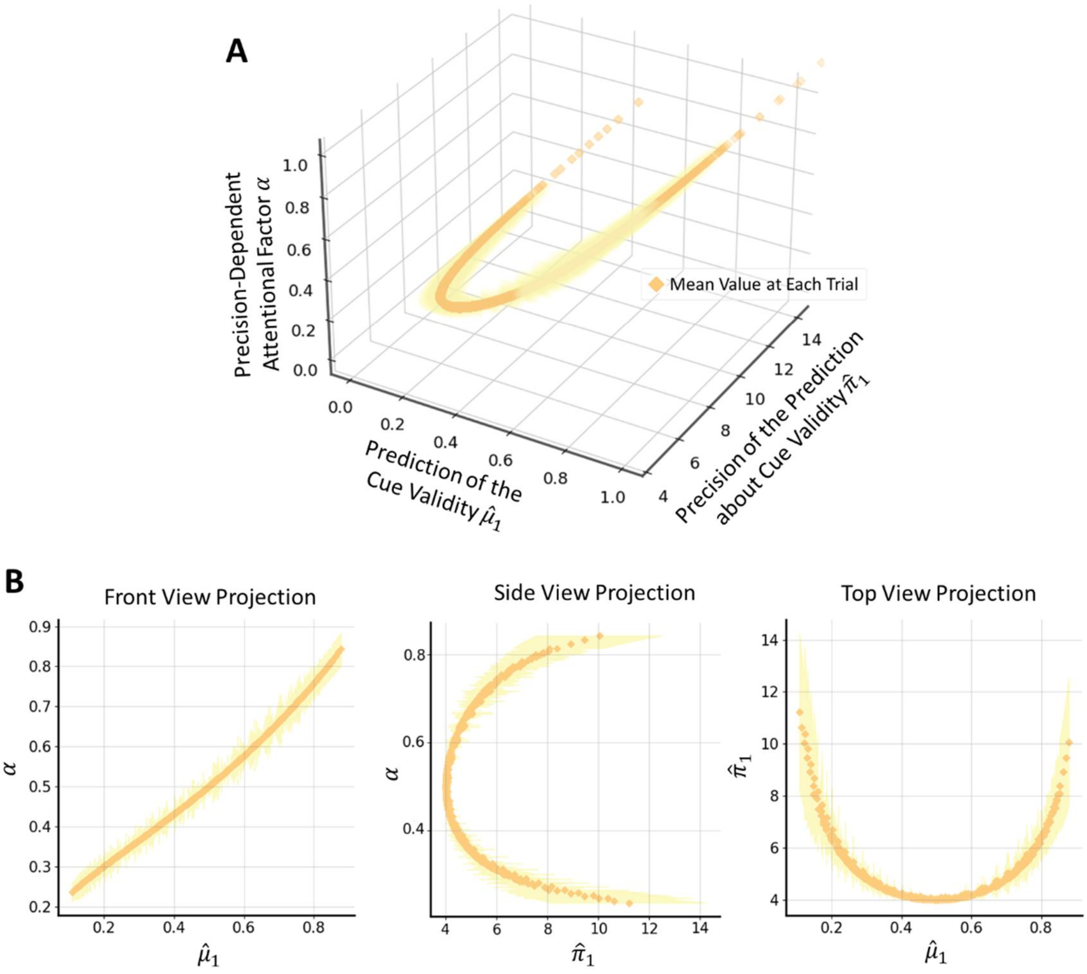


Fig. S3. (A) Relationship between $\hat{\mu}_1$ (x-axis), $\hat{\pi}_1$ (y-axis), and $\alpha$ (z-axis). Orange dots represent the mean values across all participants for each trial. (B) Projections of the three-dimensional plot in (A). Each view provides the relationship between two of the three variables by flatting (A) along one axis. Yellow shades represent standard deviation.
